# CoV2ID: Detection and Therapeutics Oligo Database for SARS-CoV-2

**DOI:** 10.1101/2020.04.19.048991

**Authors:** João Carneiro, Catarina Gomes, Cátia Couto, Filipe Pereira

## Abstract

The ability to detect the SARS-CoV-2 in a widespread epidemic is crucial for screening of carriers and for the success of quarantine efforts. Methods based on real-time reverse transcription polymerase chain reaction (RT-qPCR) and sequencing are being used for virus detection and characterization. However, RNA viruses are known for their high genetic diversity which poses a challenge for the design of efficient nucleic acid-based assays. The first SARS-CoV-2 genomic sequences already showed novel mutations, which may affect the efficiency of available screening tests leading to false-negative diagnosis or inefficient therapeutics. Here we describe the CoV2ID (http://covid.portugene.com/), a free database built to facilitate the evaluation of molecular methods for detection of SARS-CoV-2 and treatment of COVID-19. The database evaluates the available oligonucleotide sequences (PCR primers, RT-qPCR probes, etc.) considering the genetic diversity of the virus. Updated sequences alignments are used to constantly verify the theoretical efficiency of available testing methods. Detailed information on available detection protocols are also available to help laboratories implementing SARS-CoV-2 testing.

## 1. Introduction

The SARS-CoV-2 genome consists of a single, positive-stranded RNA with approximately 30 000 nucleotides. Thousands of genomic sequences are now available in public databases as the epidemic progresses. The great adaptability and infection capacity of RNA viruses depends in part from their high mutation rates.^1^ As expected, available SARS-CoV-2 genomic sequences already show a large number of polymorphisms. Many techniques use molecules that interact with the virus RNA genome or the reverse transcribed DNA, either for clinical testing, diagnosis or determination of viral loads.^2^ For example, PCR primers and RT-qPCR probes are been widely used to detect SARS-CoV-2.^3,4^ It is likely that oligonucleotides complementary to the virus RNA will be tested as possible antiviral agents.^5,6^ However, polymorphisms can be a challenge for the efficiency of available assays since they may lead to false-negative results in detection tests or inefficient therapeutics.

## 2. Materials and Methods

### 2.1 Database features

The CoV2ID database (http://covid.portugene.com/) uses java graphics and dynamic tables and works with major web browsers (e.g. Internet Explorer, Mozilla Firefox, Chrome). The database provides descriptive webpages for each oligonucleotide and a search engine to access dynamic tables with numeric data and multiple sequence alignments. A SQLite local database is used for data storage and runs on an Apache web server. The dynamic HTML pages were implemented using CGI-Perl and JavaScript and the dataset tables using the JQuery plugin DataTables v1.9.4 (http://datatables.net/). Python and Perl in-house algorithms were written and used to perform identity and pairwise calculations.

### 2.2. Oligonucleotides

The oligonucleotides were retrieved from peer reviewed publications [e.g.,^7–14^] and protocols provided by the World Health Organization (WHO). Each oligonucleotide has a specific database code (for example, *CoV2ID001*). The CoV2ID database ranks oligonucleotides using three measures of sequence conservation:

a. *Percentage of identical sites* (PIS), calculated by dividing the number of equal positions in the alignment for an oligonucleotide by its length;
b. *Percentage of identical sites in the last five nucleotides at the 3’ end of the oligonucleotide* (3’PIS) - the most critical regions for an efficient binding to the template DNA during PCR and
c. *Percentage of pairwise identity* (PPI), calculated by counting the average number of pairwise matches across the positions of the alignment, divided by the total number of pairwise comparisons.

The ‘*CoV2ID ranking score*’ considers the mean value of the three different measures (PIS, 3’PIS and PPI), as previously described.^15,16^

### 2.3 Genomic sequences

The SARS-CoV-2 ‘isolate Wuhan-Hu-1’ (NC_045512.2) was used as reference. Genomes were obtained from the NCBI GenBank (https://www.ncbi.nlm.nih.gov/genbank/sars-cov-2-seqs/) and the GISAID Initiative (https://www.gisaid.org/). The list of acknowledgments to the original source of the data available at GISAID can be found in ‘*Acknowledgments’* section of our database.

The first release of the database includes three multiple sequence alignments:

a. *CoV2ID_alig01* - SARS-CoV-2 complete genomes available at the NCBI.
b. *CoV2ID_alig02* - SARS-CoV-2 complete genomes from the GISAID with high coverage and no ambiguities or gaps.
c. *CoV2ID_alig03* - Alignment of the consensus sequence of each human coronavirus: SARS-CoV-2, HCoV-OC43, HCoV-HKU1, HCoV-NL63, HCoV-229E, MERS-CoV and SARS-CoV.

The genomes from the NCBI were aligned using an optimized version of MUSCLE running at the NCBI Variation Resource. The genomes from GISAID were aligned using the default parameters of the MAFFT version 7.^17^ The sequences can be visualized, edited and exported using the NCBI (https://www.ncbi.nlm.nih.gov/tools/sviewer/) and the Wasabi (http://wasabi2.biocenter.helsinki.fi/) tools.

## 3. Results and Discussion

The CoV2ID database currently includes 145 oligonucleotides from 21 protocols: 64 PCR primers, 57 LAMP primers, 20 probes and 4 target generation oligonucleotides. The oligonucleotides are located in the *ORF1ab*, *S*, *ORF3a*, *E*, *M* and *N* genes. The database provides an interface for browsing, filtering and downloading data from the different oligonucleotides annotated according to the SARS-CoV-2 reference genome. For each oligonucleotide, it is possible to find information on the sequence, type of technique where it was originally used, location in the reference genome, etc.

The largest multiple sequence alignment (*CoV2ID_alig02*) has 3251 complete SARS-CoV-2 genomes. The alignment has a PIS of 88.60% and a PPI of 99.96%. The NCBI alignment (currently with 106 genomes) has similar values: PIS of 62.40% and a PPI of 99.40%. These results demonstrate the existence of several mutated positions across the genome leading to relatively low percentage of identical sites. However, the level of genetic diversity is relatively low, as shown by the high percentage of pairwise identity (>99%), suggesting that most mutations only occur in a few genomes.

The database indicates which oligonucleotides bind to the most conserved regions of the SARS-CoV-2 using different measures of sequence conservation (Table 1). Several oligonucleotides have a perfect homology to all available genomes (CoV2ID score of 100%). For example, nine oligonucleotides are 100% complementary to all genomes. On the contrary, some oligonucleotides have several mismatches to SARS-CoV-2 genomes. There are six oligonucleotides with a CoV2ID score of below 60%. For example, primers *NIID_WH-1_F24381* and *NIID_WH-1_Seq_F519* have a CoV2ID score of below 50%. Previous works have already detected polymorphisms in primers and probes that may cause problems when performing the testing ^18,19^. In terms of pairs of primers, we identified 238 pairs with a CoV2ID score above 90%. For example, the pair of primers *Pasteur_nCoV_IP4-14059Fw* and *Pasteur_nCoV_IP4-14146Rv* had a CoV2ID score of 99.58%.

**Table 1.**
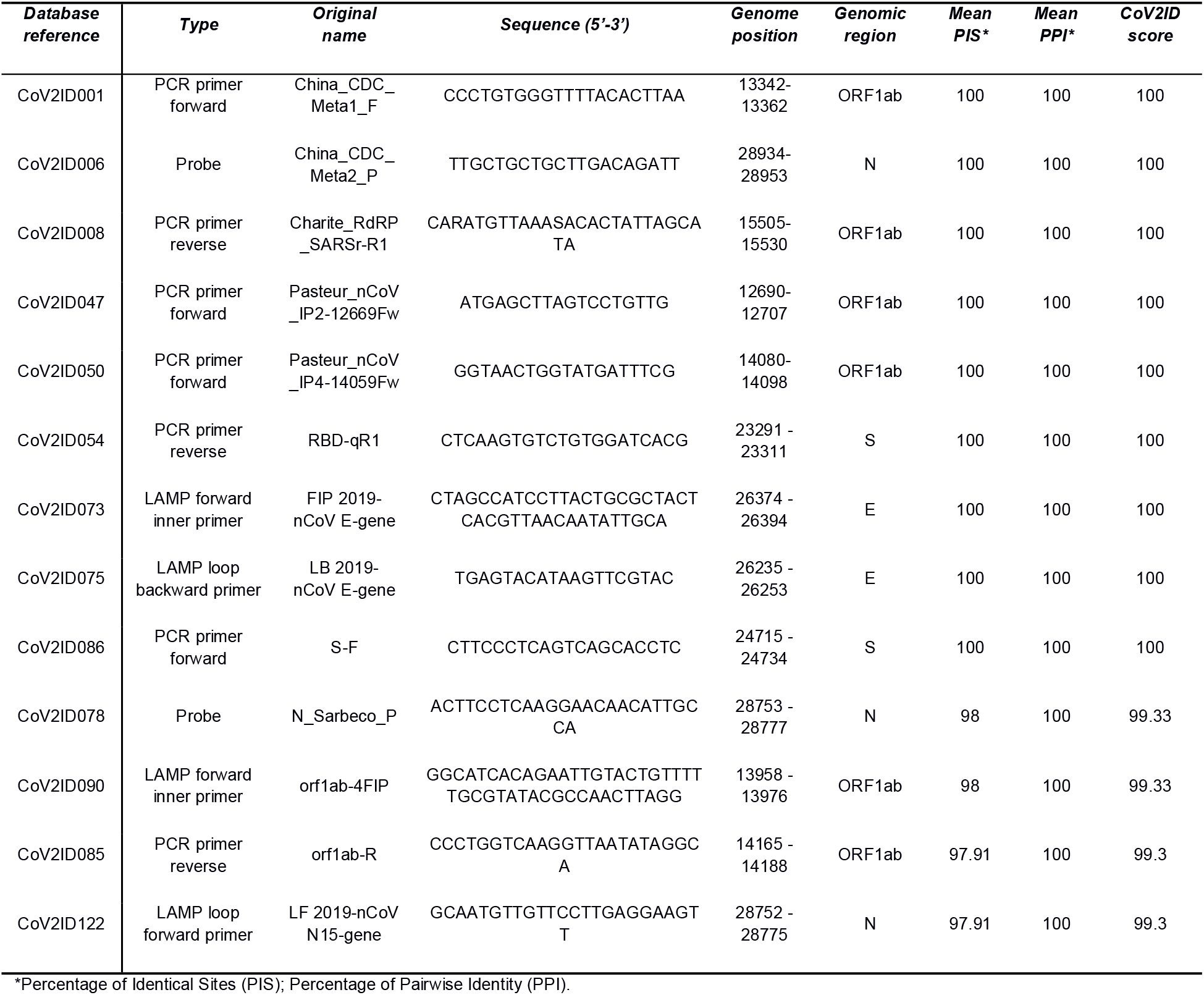
Oligonucleotides with the highest conservation score considering the multiple sequence alignments of complete SARS-CoV-2 genomes.

SARS-CoV-2 oligonucleotides with a high divergence to other strains should be preferred to avoid false positives resulting from the putative binding in non-target species. We have identified the most divergent oligonucleotides in other coronaviruses, i.e., the best ones to avoid false positives (Table 2). Fifty-six oligonucleotides have a CoV2ID score below 20% regarding other coronaviruses (*CoV2ID_alig03*), meaning they are highly divergent. On the contrary, seven oligonucleotides have a CoV2ID score above 70%. The most conserved oligonucleotide (CoV2ID102; *Chan_RdRp_gene_R*) has a CoV2ID score of 91.43%, but was designed to target all SARS-related coronaviruses ^14^, which explains its high conservation. In general, available SARS-CoV-2 oligonucleotides diverge from other human coronaviruses by several positions, and therefore are unlikely to cause false-positives.

**Table 2.**
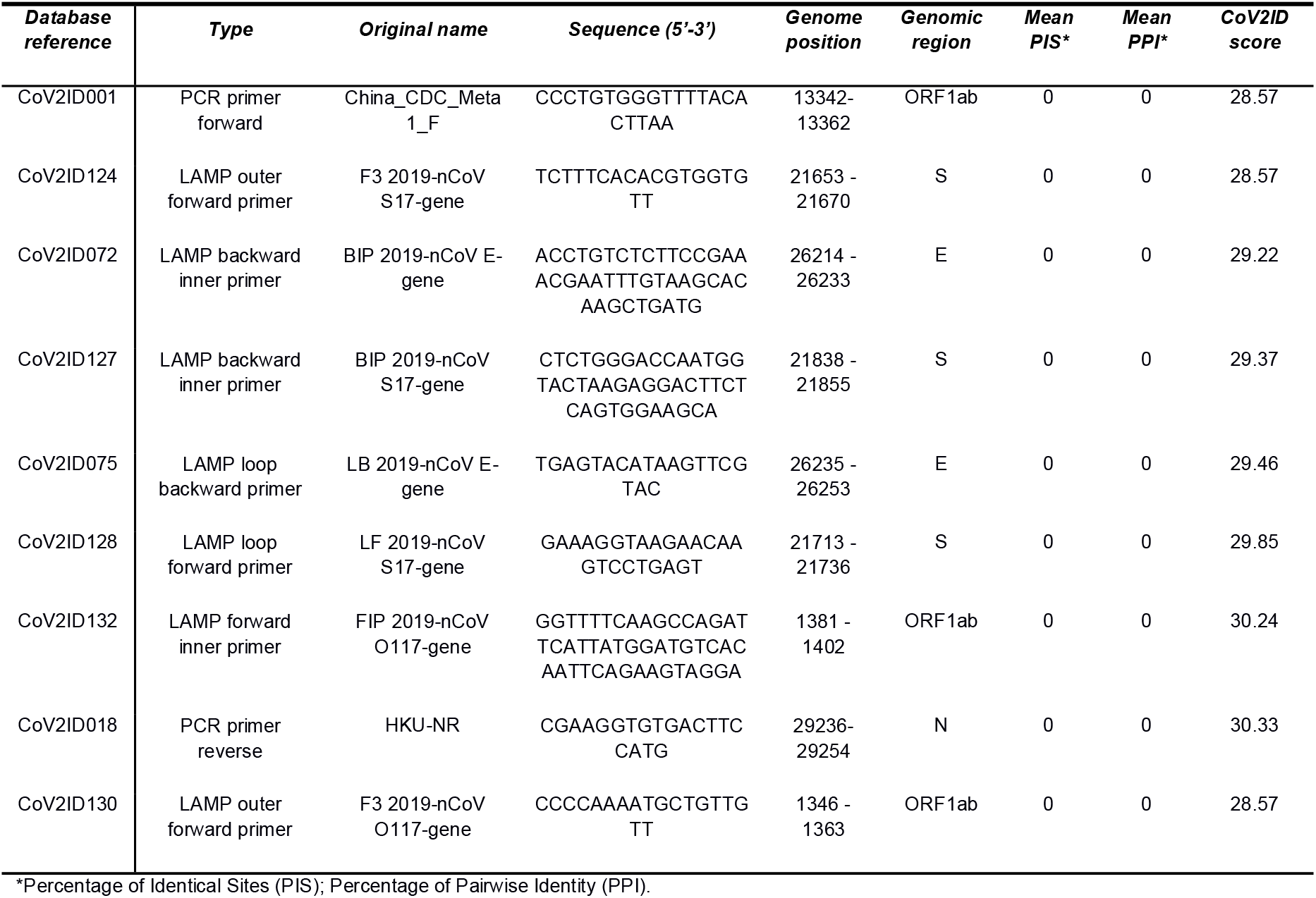
Oligonucleotides with the lowest conservation score considering the alignment of consensus sequences of all human coronavirus.

## 4. Example of use

If the aim is to choose an oligonucleotide located in a conserved genomic region, the user can navigate through the “Search” tab on the top menu bar and open the “The best oligonucleotides” tab. The table with oligonucleotides is automatically ordered by the “CoV2ID Score” column filter. The user can also access the oligonucleotide summary information by clicking in the ID hyperlink. The database can also be used to filter all columns using the search tool. If the purpose is to design a new oligonucleotide, the database section “Genome variation” should be selected in the tab on the top menu bar. The user can then visualize the PIS and PPI values in 100-nucleotide sliding windows with 50 nucleotides of overlap. The list of the most conserved genomic regions can be found in a table. In this case, the genomic regions 15901-16000 and 15951-16050 had the highest PIS value (100%) considering alignment *CoV2ID_alig02*. This section of the alignment can be visualized by clicking on the position value in the table. The user can also visualize any window of the alignment by using the ‘Show window in alignment’ box

## Acknowledgements

This research was supported by national funds through FCT - Foundation for Science and Technology within the scope of UIDB/04423/2020 and UIDP/04423/2020. J.C. also acknowledges the FCT funding for his research contract at CIIMAR, established under the transitional rule of Decree Law 57/2016, amended by Law 57/2017.

